# Controlling biases in targeted plant removal experiments

**DOI:** 10.1101/2023.08.22.554120

**Authors:** Sylvain Monteux, Gesche Blume-Werry, Konstantin Gavazov, Leah K. Kirchhoff, Eveline J. Krab, Signe Lett, Emily Pickering Pedersen, Maria Väisänen

## Abstract

1. Targeted removal experiments are a powerful tool to assess the effects of plant species or (functional) groups on ecosystem functions. However, removing plant biomass in itself can bias the observed responses. This bias is commonly addressed by waiting until ecosystem recovery, but this is inherently based on unverified proxies or anecdotal evidence. Statistical control methods are efficient, but restricted in scope by underlying assumptions.
2. We propose accounting for such biases within the experimental design, using a gradient of biomass removal controls. We demonstrate the relevance of this design by presenting i) conceptual examples of suspected biases and ii) how to observe and control for these biases.
3. Using data from a mycorrhizal association-based removal experiment we show that ignoring biomass removal biases (including by assuming ecosystem recovery) can lead to incorrect, or even contrary conclusions (e.g., false positive and false negative). Our gradient design can prevent such incorrect interpretations, whether aboveground biomass has fully recovered or not.
4. Our approach provides more objective and quantitative insights, independently assessed for each variable, than using a proxy to assume ecosystem recovery. Our approach circumvents the strict statistical assumptions of e.g. ANCOVA and thus offers greater flexibility in data analysis.

## Introduction

Plants are terrestrial organisms that govern over soil physical structure, pedogenesis, hydrology and biogeochemical cycling. Changes in vegetation composition may therefore have far-reaching consequences for ecosystem services. Plant removal experiments are a widespread method to assess the effects of vegetation changes on ecosystem functioning (Morais & Cianciaruso, 2014). By allowing to assess ecosystem responses from soil molecular and microbial scale to aboveground vegetation community responses, plant removal experiments have produced valuable insights on the ecological role of given plant functional groups or plant species, however to the best of our knowledge they have seldom targeted plants based on their mycorrhizal associations.

Removing plant groups, however, comes at the price of strong impacts on the ecosystem, caused notably by severing the energy flow through rhizodeposition, by altering shading and evapotranspiration, or by producing root necromass (Aarssen & Epp, 1990; Campbell *et al*., 1991; McLellan *et al*., 1995; Díaz *et al*., 2003; Mikola *et al*., 2014; De Long *et al*., 2016). Studying these impacts is essential when focusing on plant competition, facilitation or nursery effects, which constitute the basis of the wide field of “neighbor removal” experiments. On the contrary, targeted plant removal experiments aim at understanding the specific role of a given group of plants: it is therefore necessary to untangle the effects of removing biomass in general (hereafter, “non-specific biomass removal”) from the effects of removing biomass of the targeted plants, *i*.*e*., of removing *a* plant compared to removing *this* plant. This is especially problematic in systems with an uneven abundance of the target groups, where the large difference in amount of biomass removed between different groups may conceal putative effects of targeted removal. Here, we go through the scenarios that can arise from such biases and the existing methods of control, before proposing to combine experimental and statistical approaches to correct for these unwanted biases in targeted plant removal experiments.

Most targeted removal studies today assume an ideal “steady-state” scenario, where, after an adequate duration, the system has entirely recovered from initial disturbance. In this case, no matter how much biomass was initially removed, the disturbance is assumed to no longer affects the system. We hereafter refer to this case as *scenario (a)* (Fig. 1a). This scenario depicts a situation where non-specific biomass removal does not induce a measurable effect on the response variable and therefore any observed targeted removal treatment effect is indeed a “true” effect. This means that the apparent effect of targeted removal is not biased by effect of the biomass removal itself and does not therefore need further control.

**Figure 1:**
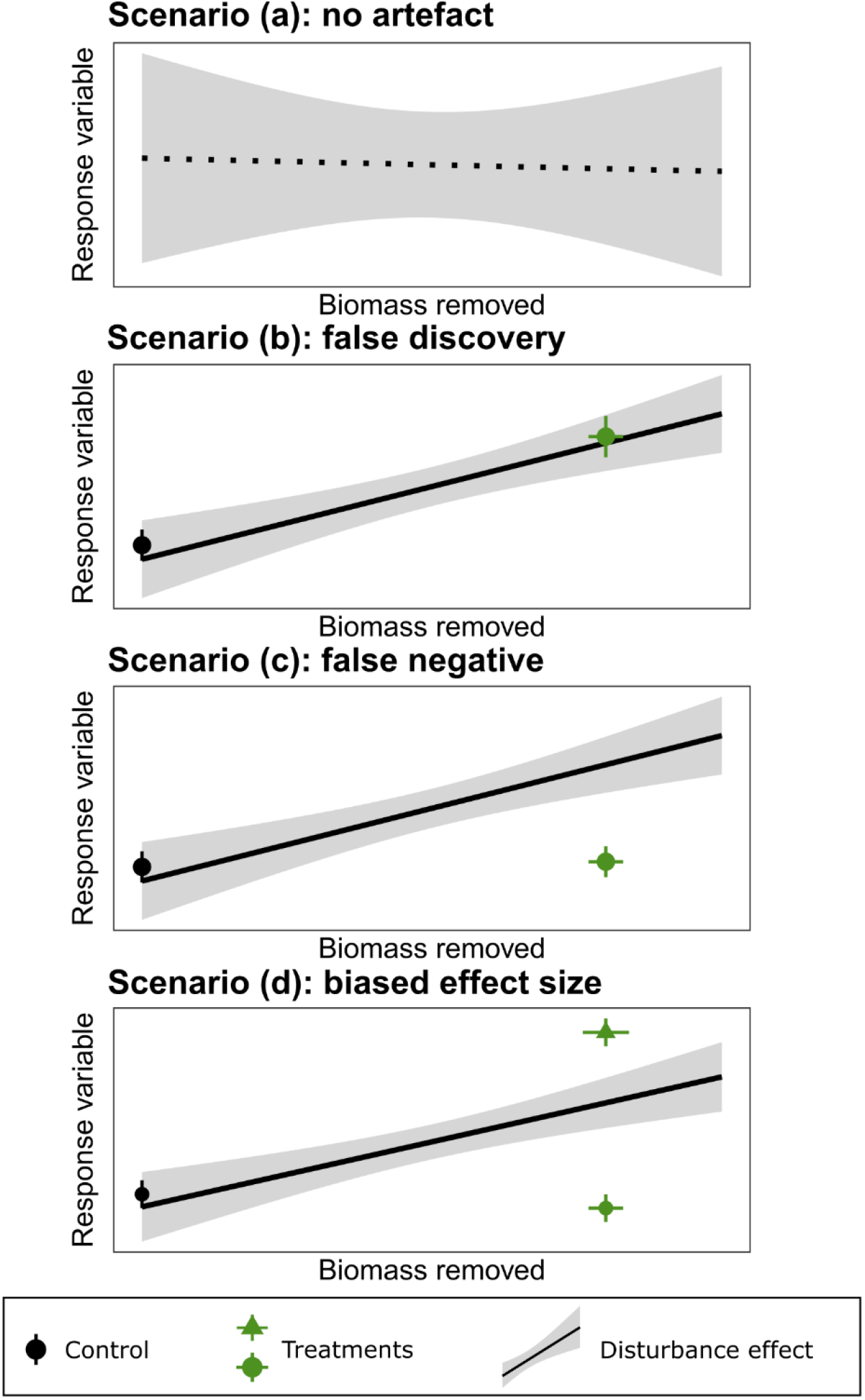
Artefacts associated with non-specific biomass removal. When non-specific biomass removed does not significantly relate to the response variable (dotted line, **a**), data obtained after targeted removal can be analyzed without adjustment. Ignoring a significant relationship between non-specific biomass removal and the response variable (solid lines, **b-d**) can lead to erroneously attributing an effect to a targeted removal treatment when it is in fact due to removed biomass itself **(b**), or, conversely, disregarding as a non-significant difference a strong treatment effect that negates that of removed biomass (**c)**. Misestimation of effect sizes (**d**) can also occur, resulting in over-(green triangle) and under-estimates (green circle).

In contrast to *scenario (a)*, non-specific biomass removal may associate significantly with a response variable (*e*.*g*., response turns more positive/negative as more biomass has been removed), and under these conditions several types of biases can arise:

- One such bias is referred to as “false discovery” in *scenario (b)*: the treatment effect for a response variable may initially appear as strongly significant (Fig. 1b), but may in fact be entirely caused by non-specific removal effect. In other words, if the effects of removing similar amounts of non-specific and targeted biomass are indistinguishable, then there is no ground to attribute the observed effect to targeted biomass removal.
- Conversely, a targeted removal treatment may initially appear to have no significant effect on a given response variable if the treatment and its control do not differ from each other (Fig. 1c), while, at the same time, the non-specific removal associates with the response variable. The non-specific removal effect may counteract the effect of targeted removal, resulting in no apparent differences between the targeted removal treatment and its control. This overlap can lead to dismissing a strongly significant effect of targeted removal as insignificant, due to the confounding effect of non-specific removal, we therefore refer to this scenario as the “false negative” *scenario (c)*.
- Aside from cases of false discovery and false negative, any significant relationship between a response variable and the disturbance can be expected to bias – overstate or downsize – the observed effect size of treatment effects (*scenario (d)*, Fig. 1d).

While the biases caused by sheer non-specific removal of biomass may not be controlled, they are often considered negligible after allowing the system to recover, and its recovery, in turn, is estimated through various proxies. One proxy to estimate recovery after disturbance is aboveground vegetation, through its cover or biomass (e.g., Wardle & Zackrisson, 2005; Gundale *et al*., 2010), which may address some plant-related aspects of the disturbance brought to the ecosystem, such as the interruption of energy flow belowground or changes in micrometeorological conditions. However, it always remains unclear whether these proxies are adequate for other elements of the ecosystem (e.g., belowground or non-plant), and assessing this for multiple variables is impractical or outright impossible. Furthermore, whether recovery proxies are suited to systems where recovery may require several decades, such as arctic ecosystems (Cannone *et al*., 2010; Mikola *et al*., 2014), is also questionable. Finally, competition or facilitation may result in a steady-state aboveground biomass that is lower or higher than prior to removal, which may question the relevance of this proxy. A few commendable studies attempt to validate the assumption of ecosystem recovery (e.g., Wardle & Zackrisson, 2005), but this is rarely the case. Thus, it is common practice in this field to arbitrarily consider that the ecosystem has recovered without any quantitative support.

Another approach to address disturbance-induced biases, even on shorter time scales, is to use statistical control with methods such as analyses of covariance (ANCOVA, e.g., Wardle & Zackrisson, 2005) with removed biomass as a covariate. However, this is not applicable in all experimental designs. Indeed, ANCOVA strictly requires a linear relationship between the dependent variable and its covariate and is thus not relevant for many data requiring different analyses, such as generalized linear models (e.g., abundance data). Even if the linearity criterion is satisfied, an underlying assumption of ANCOVA is that the response-covariate relationship is identical across levels of the independent variable, an assumption that is complicated when target plant groups abundances widely differ, and which simply cannot be met while including a control group without any removal (therefore without any slope). In such cases, other statistical control tools may be used such as multiple regressions (e.g., Bret-Harte *et al*., 2004) or structural equation models, but these tools also impose their own statistical constraints such as linearity assumptions or high replication. Studies approaching the problem through experimental design are promising but represent only a fraction of targeted removal studies, even among those that take into account biases due to biomass removal (e.g., Symstad & Tilman, 2001; Rewcastle *et al*., 2022).

Here we introduce an experimental and analytical design to account for the effects of non-specific plant biomass removal based on a gradient of non-specific removal. We validate the approach with data from an experiment aiming to study above- and belowground linkages under changing vegetation at the forest-tundra ecotone in sub-arctic Sweden. Strong changes in abundance of plant functional groups are expected in the Arctic with climate change (Elmendorf *et al*., 2012; Myers-Smith *et al*., 2015; Mekonnen *et al*., 2021), accompanied by changes in mycorrhizal associations which might affect the large carbon pool in arctic soils (Clemmensen *et al*., 2021). Our experiment thus targets plants for removal based on their mycorrhizal associations. We 1) conceptually present the different biases that can arise from not accounting for biomass removal effects, 2) provide examples of above- and belowground variables from our field experiment (β-xylosidase activity, *Betula nana* growth and leaf δ^15^N) where not accounting for these biases would lead to misestimating or wrongly interpreting observed findings, and 3) indicate how we adjusted the data to control biases.

## Methods

### Study area and site

The experimental site is located in sub-arctic northern Sweden south-east of Abisko on a WNW-facing slope approximately 515 m above sea level. The local climate is continental sub-arctic (30-year mean annual air temperature 0.18 °C, displaying an increasing trend; Callaghan *et al*., 2010; Abisko Scientific Research Station, 2023), with mild oceanic influences, while a rain shadow maintains low precipitation (300-400 mm yr^-1^). Regional vegetation is dominated by mountain birch forest (*Betula pubescens* Ehrh. ssp *czerepanovii* (Orlova) Hämet Ahti.) at low elevation, transitioning at the treeline ecotone (500 – 600 m a.s.l.) into ericaceous-dominated heath interspersed with graminoid-rich meadow tundra.

Our study site is located at the birch forest-tundra ecotone and underlain by base-rich schist (Jämtlandian Caledonian orogen of the Offerdal and Särv Nappes; Sveriges geologiska undersöknings, 2023). The vegetation at the site is a heath tundra dominated by *Empetrum nigrum* ssp *hermaphroditum* L., *Vaccinium uliginosum* L., *Cassiope tetragona* D. Don, *Betula nana* L. and multiple species and hybrids of *Salix* spp.. *Andromeda polifolia* L. and *V. vitis-idaea* L. are also ubiquitous but due to their low stature do not form a substantial part of the canopy. Due to the wind exposure, snow cover thickness in the winter is generally less than 20 cm (personal observations) and is continuous from late October-early November until spring melt in May-June. Consequently, the plant growing season is constrained between June and early September for the aboveground parts, and until October for roots (Blume-Werry *et al*., 2016). No permafrost is found at the study site.

### Experimental design

We tested the relevance of our suggested gradient design by assessing the effects of targeted and non-specific plant biomass removal, within a removal experiment targeting plants based on their mycorrhizal associations. To this end, the experiment was designed to a) selectively remove plants with either ectomycorrhizal, ericoid mycorrhizal or both associations and b) account for the effects of non-specific biomass removal such as disturbance or decreased rhizodeposition. We established over growing seasons 2018 and 2019 five spatially-replicated blocks over an area of 4 ha, the distance between adjacent blocks being 20–100 m. Each block contained three mycorrhizal removal treatment plots, three non-specific biomass removal gradient plots and one undisturbed control, and the size of each plot was 2×2 m (Fig. 2). To ensure the absence of clonal plant growth or mycorrhizal hyphae from the area outside of the plots, the borders of the plots were trenched with a breadknife and a shovel down to the bedrock or to 30 cm depth two to three times over the course of each growing season.

**Figure 2:**
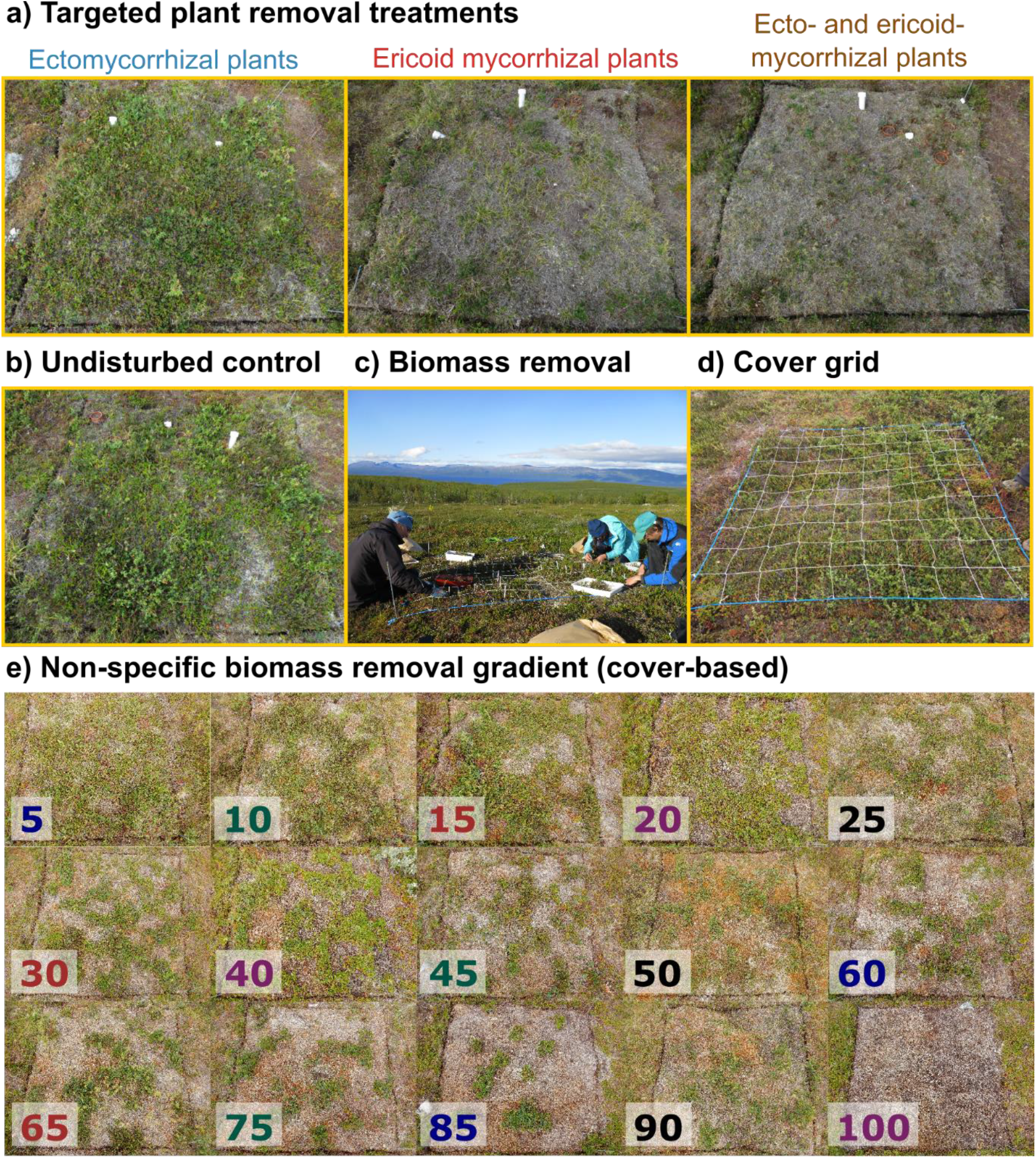
Experimental design. **(a-b)** The core experiment consists in a factorial specific removal of ectomycorrhizal and ericoid mycorrhizal plant removal, combined with an undisturbed control, (**c**) Example of a non-specific biomass removal gradient plot before removal: a nylon grid is overlain on the plot where each grid cell represents 1% of the cover (20 x 20 cm), (**d**) Biomass removal in a non-specific biomass removal gradient plot, (**e**) The different levels of removal used in the biomass removal gradient. Each of the five spatial blocks contains a control, three treatment plots and three nonspecific biomass removal gradient plots, to quantify and adjust for disturbance effects induced by removing plant biomass. Gradient plots within each block share the same font color and represent each of the low (5-25), medium (30-60) and high (65-100) removal categories.

### Mycorrhizal removal

Mycorrhizal removal treatments consisted of selectively removing ectomycorrhizal (hereafter referred to as -ECM), ericoid mycorrhizal (hereafter -ERM) or both ectomycorrhizal and ericoid mycorrhizal plants (hereafter -ECM/-ERM). The mycorrhizal associations of all plant species found within the site were assigned based on existing databases (Harley & Harley, 1987; Akhmetzhanova *et al*., 2012), and further confirmed by microscopy observations on roots from 10 individuals of each species sampled outside of the plots in July 2019. Ectomycorrhizal plant species targeted by removal were *B. nana, Salix* spp., *Bistorta vivipara* L., and *Dryas octopetala* L., and amounted to *c*. 13% of vegetation cover, while ericoid mycorrhizal plants were *E. nigrum, V. uliginosum, V. vitis-idaea, C. tetragona, A. polifolia*, and *Rhododendron lapponicum* L. and amounted to *c*. 85% of vegetation cover. *Arctous alpina* (L.) with *c*. 20% cover is the main arbutoid mycorrhizal plant, facultative arbuscular mycorrhizal plants include *Nardus stricta* L. and *Festuca ovina* L. but no arbuscular mycorrhiza were observed, non-mycorrhizal plants include e.g. *Carex* spp. and *Pedicularis lapponica* L., and most plant species at the site harboured dark septate endophytes which may provide mycorrhizal-like functions (Newsham *et al*., 2009).

### Biomass removal gradient

To control the biases induced by biomass removal in the mycorrhizal removal treatment plots, we established a gradient of non-specific biomass removal. To this end, we removed a given percentage of the biomass within three removal gradient plots in each block (Fig. 2). The amount of biomass removed in each gradient removal plot was allocated randomly, with the constraint that each spatial block should contain one of each low (5–25 %), medium (30–60 %), and high (65–100 %) levels of biomass removal intensity. To remove a given amount of biomass, a mesh with 100 20×20 cm squares was overlain on the plot, each square representing 1 % of the surface. The adequacy of area as a proxy for biomass removed is shown in Supplementary Figure S1 (Pearson’s r = 0.95; 95%CI 0.86 -0.98; df = 13; *P* < 0.001). The 20×20 cm squares where vegetation was removed were allocated randomly, with the constraint that a similar number of squares were present in each of the four 1 m^2^ quarters constituting the plot.

### Vegetation removal

Woody biomass was clipped with scissors just below the soil surface to minimize disruption of the soil structure (McLellan *et al*., 1995) that would result from pulling out roots of woody species (e.g., *B. nana, E. nigrum, V. uliginosum*), while non-woody vegetation or thin woody stems were either clipped or plucked out without disturbing the soil surface. Vegetation was removed in an identical way in targeted removal and non-specific biomass removal plots. Removed biomass was air-dried at 40 °C for up to 14 days, then weighed. The removal was repeated yearly during the growing seasons of 2018 to 2021.

### Response variables

#### β-xylosidase activity

Microbial extracellular enzymatic activity of β-xylosidase was measured from soils collected 14–15 July 2021 from the humus (top ∼4 cm) horizon using an apple corer (⌀ 1.5 cm). Eight cores (two in each 1 m^2^ quarter of a plot) were pooled to form a composite sample stored at 4 °C and homogenized through a 2 mm sieve within the day. β-xylosidase activity was measured from these fresh soils within three days with the Soil Enzymatic Activity Reader (SEAR; Digit Soil, Zürich, Switzerland). The measurement is based on the reaction of soil enzymes with enzyme-specific fluorogenic substrate namely 4-Methylumbelliferyl-β-D-xylopyranoside (MUX). In each measurement Methylumbelliferyl (MUF) standards were included to calibrate the measurement.

#### Plant growth, N and δ^15^N content

The growth of an ectomycorrhizal plant, dwarf birch (*B. nana)*, was monitored over the course of one growing season (June-August 2021) by measuring shoot elongation. Shoots from four individual stems per each control or -ERM plots (n = 40, *B. nana* absent in -ECM and -ECM/-ERM plots) and per each biomass removal gradient plots excluding the 100% removal (n = 56) were marked with plastic cable ties in late June. Their growth was measured as the change in distance from the beginning of the marked branch segment to the branch tip, between July 1^st^ and August 25^th^. The growth from these four pseudo replicates was then averaged within each plot.

In July 2021, a total of 10–20 healthy-looking, fully expanded, and green leaves were collected from one *B. nana* individual per plot in the control (n = 5), -ERM (n = 5), and biomass removal gradient (n = 14, no data from the 100% removal plot) treatments. The leaves were oven-dried at 60 °C for 48 h, then ball mill ground and two milligrams were weighed into Sn capsules. These were analysed for nitrogen content and isotopic ^15^N/^14^N ratio on an Element Analyzer Euro EA3000 coupled to a Delta V Advanced Isotope Ratio Mass Spectrometer (EA-IRMS) at the Swiss Federal Institute for Forest, Snow and Landscape Research WSL. Stable isotope ^15^N abundance is reported in permil using the δ notation and the ^15^N / ^14^N ratio of air as a standard: δ^15^*N* = (*R*_*sample*_/*R*_*standard*_) × 1000, with a measurement error of 0.06 permil (32 repeats of NIST-Standard 1547 Peach Leaves 996).

#### Statistical analyses

All statistical analyses were carried out in the R environment (4.1.3; R Core Team, 2023), using the package lmerTest (3.1-3; Kuznetsova *et al*., 2017), lme4 (1.1-33; Bates *et al*., 2015), and the ggplot2 graphical display ecosystem (3.4.0; Wickham *et al*., 2016), with additional libraries sciplot (1.2; Morales *et al*., 2020), tidyverse (1.3.2; Wickham *et al*., 2019), patchwork (1.1.2; Pedersen, 2022) and ggside (0.2.1; Landis, 2022).

Our approach can be summarized by a simple algorithm: if we observed a significant association between a response variable and the amount of biomass removed in the gradient plots, we adjusted the response variable in treatment plots by subtracting the predicted effect of biomass removal prior to analysis; otherwise, we analysed the observed data without adjustment (Fig. 3).

**Figure 3:**
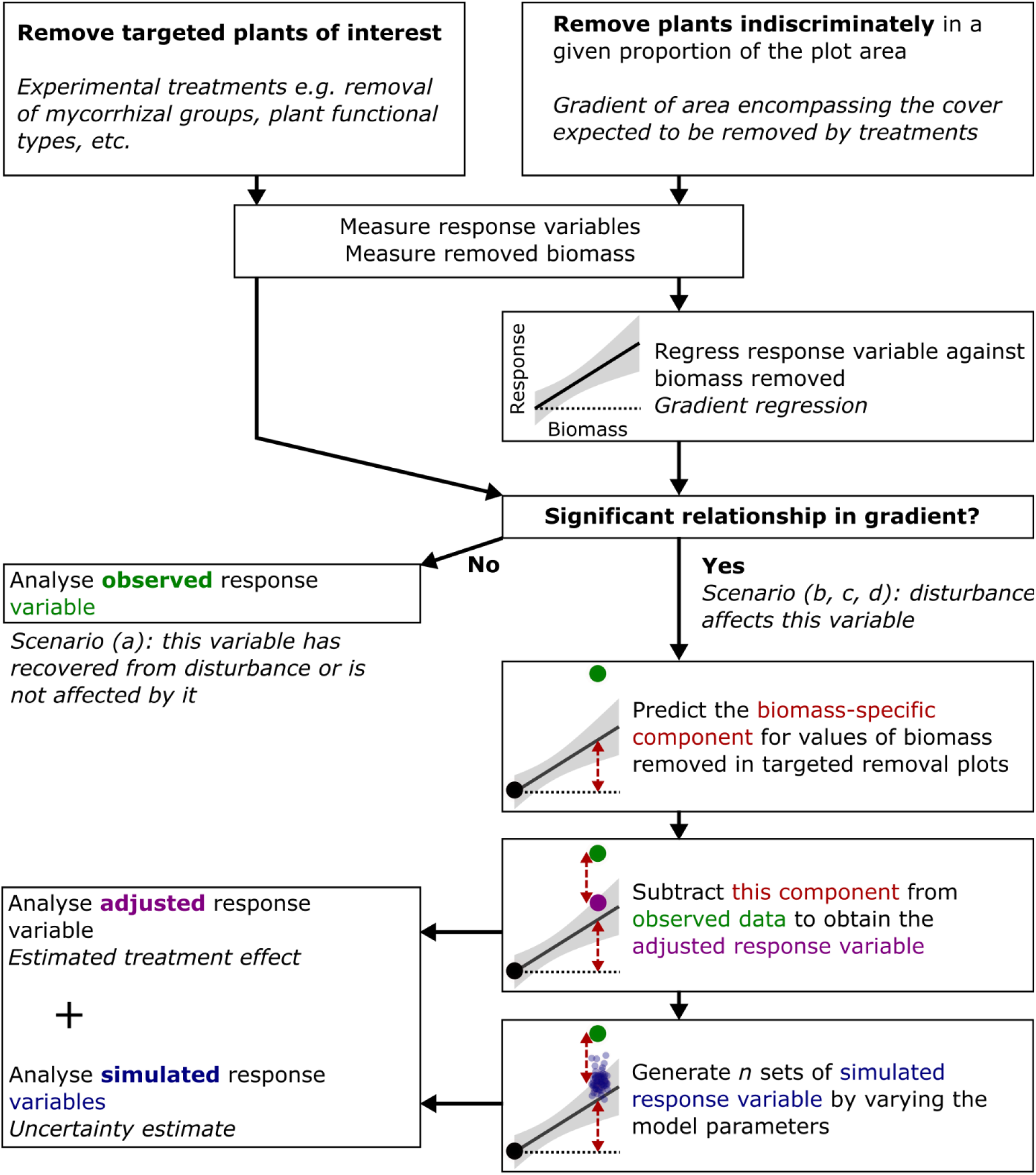
Algorithm for adjustment and analysis of observed data in the suggested experimental design.

Within the biomass removal gradient, we used linear mixed regression models to test whether the non-specific biomass removal affected the response variable (soil β-xylosidase activity, growth and leaf δ^15^N of *B. nana*). We therefore regressed each response variable against the sum of biomass removed across all years prior to the measurement, using block as a random factor. We further refer to these models as the “gradient regressions” to ease wording. If the slope of the gradient regression did not significantly (*i*.*e*., *P* > 0.05) differ from 0 (*scenario (a)*, β-xylosidase activity, Fig. 4), non-specific biomass removal had no detectable effect on the response variable in question and, consequently, we considered there was no bias due to biomass removal. In this case, we analysed the data from the mycorrhizal removal treatment plots with a two-way ANOVA on linear mixed effects models, using -ERM and -ECM treatments as fixed factors and block as a random factor.

**Figure 4:**
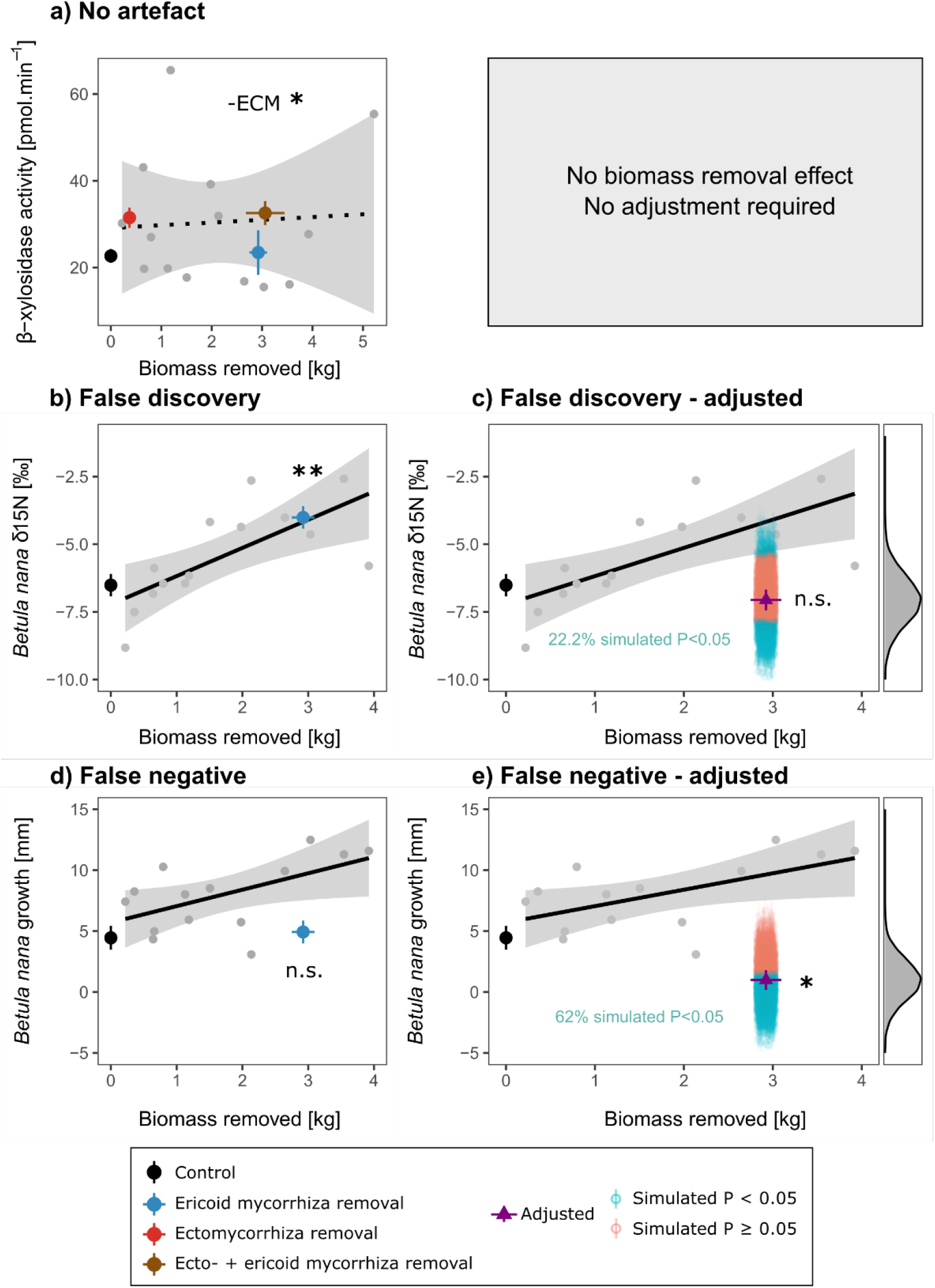
Artefacts associated with non-specific biomass removal and adjustment using a disturbance gradient. The left-hand column presents the observed data, with the means +/-SE (n = 5) of control and treatments (large coloured symbols), as well as the individual points from the non-specific biomass removal gradient (grey dots), the corresponding gradient regression (black line, solid when regression *P* < 0.05, otherwise dotted) and its 95% confidence interval (grey shaded area). The right-hand side presents the adjusted and simulated data, except for (**a**) where no adjustment was necessary: purple triangle represents the mean +/-SE (n = 5) of the adjusted response variable. The blue and red transparent symbols each represent the mean +/-SE of one of the 5000 simulated datasets, respectively when the *P* value of the ANOVA test for the -ECM treatment effect is below or above 0.05. The simulated points have the same value on the x-axis and are only spread out for readability. The insets on the right-hand side represent the density distribution of simulated means. *: *P* < 0.05; **: *P* < 0.01; n.s.: *P* > 0.05.

When the slope of the gradient regression significantly differed from 0, we considered that the non-specific biomass removal affected the response variable in question (*scenarios (b-d), B. nana* growth and δ^15^N, Fig. 4) and, consequently, induced bias. We extracted the gradient regression slope to account for the confounding role of non-specific biomass removal on the response variable. We adjusted the response variable by subtracting the biomass-dependent component, predicted by the gradient regression model, from it, and then used two-way ANOVAs on the adjusted response variable, using -ERM and -ECM treatments as fixed factors and block as a random factor.

For all variables in the gradient plots, as well as for *B. nana* growth in the treatment plots, singular model fits resulted in identical intercept values across blocks. For the sake of consistency, we carried on the analyses on these models anyway, which essentially comes down to using linear models without random factors in cases where this occurred.

#### Monte Carlo simulation

Subtracting the predicted biomass-dependent component from the response variable (measured in targeted removal treatments) provides an adjusted response variable that corrects for the location effect of the non-specific biomass removal, but not its dispersion. In other words, the solid line in Fig. 1 is taken into account by this approach, but not its associated confidence interval. Thus, while the trend of the biomass-response variable relationship is appropriately accounted for, its variability is not, even though this variability can differ between response variables and over different amounts of biomass removed. We thus assessed to what extent the variability associated with the gradient regression model could affect our conclusions by conducting Monte Carlo simulations of the response variable adjustment. For variables where an adjustment was necessary (Fig. 4b and 4d), we simulated 5000 slope values following a normal distribution centred on the slope estimate and dispersed using the slope standard error, as identified by the gradient regression model. We used these simulated slopes to predict the values of the adjusted response variables (5000 sets of 5 samples), and then tested whether these sets differed from the observed values in the Controls, using ANOVA on linear mixed models as described above. The distribution of the means (±SE, n = 5) of these 5000 sets for each simulated response variable is shown in Fig. 4c and 4e, as well as the proportion of those sets which significantly differed from the Controls (ANOVA *P* < 0.05). This provides a measure of the uncertainty associated with our adjustment process, allowing more confidence in attributing any discrepancy between the observed and adjusted variable to chance or to an actual artefact.

#### Non-linear approach

For the sake of an example of a non-linear approach, we also analysed *B. nana* growth using generalized linear mixed models. Shoot elongation is strictly positive, we therefore used a Gamma distribution for this example to ensure positive numbers and used an inverse link, which resulted in marginally lower AIC than log or identity links. A generalized linear mixed model (GLMM) could be fitted for the non-specific biomass removal gradient plots, which fit was sensibly as good as the linear regression above. The adjustment process on *B. nana* growth data in the treatment plots resulted in some negative values, which could not be analysed downstream by GLMM and were thus converted to a small positive number (10^-5^) prior to analysis. After adjustment, the singular fit of the GLMM led us to analyse the adjusted response variable using a GLM without including block as a random factor. A simple uncertainty assessment as described above would have been inadequate in this case, and carrying out a more accurate uncertainty assessment was beyond the scope of this example.

## Results

We considered different conceptual scenarios in which biomass removal – or disturbance – could lead to incorrect interpretations of observed findings (Fig. 1). We then identified variables within our data that correspond to these scenarios (Fig. 4, left column), and present how we did or did not adjust these variables to illustrate the relevance of our approach (Fig. 4, right column).

### No biomass removal artefact

The activity of β-xylosidase corresponds to the “no artefact” *scenario (a)* presented in Fig. 1a: there was no significant (*F*_1,8.3_ = 0.553, *P* = 0.478, Fig. 4a) relationship between β-xylosidase activity and non-specific biomass removal (Fig. 4a). Thus, there was no bias due to non-specific biomass removal and we could identify a significant increase in β-xylosidase activity with -ECM treatment in comparison to the control (2-way ANOVA *F*_1,14_ = 5.960, *P* = 0.029). There was no difference between -ERM and control treatments (*F*_1,14_ = 0.064, *P* = 0.804) and there was no significant interaction between -ECM and -ERM treatments (*F*_1,14_ = 0.001, *P* = 0.972).

### False discovery

The δ^15^N of *Betula nana* leaves corresponds to *scenario (b)*: there was a significant relationship between *B. nana* δ^15^N content and removed biomass in the gradient regression (*F*_1,12_ = 11.551, *P* = 0.005). There was also a significant effect of the -ERM treatment (*F*_*1,4*_ = 44.673, *P* = 0.003), and not accounting for biomass removal could lead to a false discovery. Indeed, when the non-specific biomass removal effect was subtracted from the observed data (purple triangle in Fig. 4c), we could no longer observe any difference between the control and the -ERM treatment (*F*_*1,4*_ = 1.515, *P* = 0.286). The distribution of simulated values revealed that only 22.2% of the 5000 simulations differed significantly (*P* < 0.05) from the controls after adjusting for the removed biomass effect, in both the lower and upper ends of the distribution. Once adjusted for the non-specific biomass removal effect, *B. nana* δ^15^N in -ERM treatment plots thus became virtually identical to that in the undisturbed control.

This is strong evidence that the significant treatment effect observed initially was a false discovery, in other words an artefact owing to plant biomass removal rather than the removal of ericoid mycorrhizal plants specifically.

### False negative

The growth of *B. nana* illustrates *scenario (c)*: growth was related to biomass removed in the gradient regression (*F*_1,12_ = 5.495, *P* = 0.037). We initially could not see a difference in growth between the control and -ERM treatment (*F*_*1,4*_ = 0.126, *P* = 0.732), but the non-specific biomass removal effect could counteract the effect of ericoid plant removal. Indeed, when the non-specific biomass removal effect was accounted for, ericoid-removal revealed a significant decrease in the growth of *B. nana* compared to the controls (*F*_*1,4*_ = 7.373, *P* = 0.026). This strong decrease (-78%) was significant in the majority (62%) of the simulations we conducted to account for the confidence interval around the gradient regression (Fig. 4e). The response variable adjustment produced some negative growth values, which are not expected to be observed in nature, we therefore re-analysed the data by converting those negative values to zero, for sensibly identical findings (-70% growth in -ERM compared to control, *F*_*1,4*_ = 7.226, *P* = 0.028, 61.6% of simulations differing from Control, Supplementary Figure S2). Therefore, the non-significant effect observed initially was likely a “false negative”, *i*.*e*. a failure to detect an existing significant effect due to the underlying non-specific biomass removal effect.

Analysing *B. nana* growth with GL(M)M, yielded sensibly identical results. There was a significant effect of non-specific biomass removal in the gradient model (*Χ*^*2*^ = 5.369, *P* = 0.021, df = 1), but no effect of ericoid-removal in the observed data (*Χ*^*2*^ = 0.139, *P* = 0.709, df = 1). After adjusting for the effect of removing non-specific biomass, an effect of ericoid-removal was noticeable (*Χ*^*2*^ = 6.581, *P* = 0.010, df = 1) of similar magnitude as with the linear approach (-83%, Supplementary Figure S2).

## Discussion

Controlling for biases due to non-specific biomass removal through experimental design has been attempted in different ways, occasionally through the use of positive controls (e.g. disturbed without biomass removal; Bret-Harte *et al*., 2004) and very rarely by removing non-specific biomass (Symstad & Tilman, 2001; Mariotte *et al*., 2013; Rewcastle *et al*., 2022). In an overwhelming majority of cases, however, this is done by waiting for the system to recover from disturbance and reach a new steady-state. It is not uncommon to read that the system is considered to have recovered from disturbance, without basing this on any measurement. This is obviously not ideal yet entirely understandable, considering that waiting for full recovery can be unpractical, especially in systems where this might require longer than an entire career (e.g., 50 years; Cannone *et al*., 2010). Using a proxy for ecosystem recovery, such as aboveground plant biomass or vegetation cover, likely provides more reliable estimates of real recovery than subjective assessments not grounded in data. However, this may be inadequate for different variables, for instance a decoupling between above- and belowground plant biomass could persist, as could an altered belowground system when large amounts of un-decomposed root biomass are present. Aboveground vegetation may also be inadequate as the new steady-state could have more or less vegetation than prior to removal, as discussed above.

Our approach circumvents such pitfalls, by controlling for disturbance effects no matter whether the system has fully recovered or not: before full recovery, our gradient approach allows to subtract the effect of non-specific removal from that of targeted removal, as detailed above. Once the system has fully recovered, by definition the relationships between response variables and non-specific biomass removal will no longer be significant. Indeed, if the system has recovered from disturbance induced by removing biomass, response variables should no longer associate with the amount of removed non-specific biomass, therefore supporting with data that adjustments are not (or no longer) necessary.

Incidentally, this also allows for detection of delayed responses of particular variables to the initial disturbance: element pools with small net fluxes, such as soil organic carbon content, or the biomass of a saprotroph specialized in very recalcitrant necromass could remain unaffected by the treatments and associated disturbance for several years before responding. Such a delayed response may still be driven mostly by disturbance, but could become visible even after the recovery proxy (*e*.*g*., plant biomass) suggests that the system has recovered. This could lead to the misleading conclusion that a particular treatment affected a variable, when it was in fact driven by disturbance. In this case, however, the response variable would associate not only with the targeted removal treatment, but also with non-specific removal biomass. Importantly, our design would allow to detect such associations and therefore identify delayed responses to initial disturbance as such.

Statistical control of initial disturbance is a less-commonly used approach but can help to rule out some artefacts of plant removal. Analyses of covariance (ANCOVA) or multiple linear regressions are typically used in this context to separate information explained by the covariate (the removal of biomass itself) from that explained by the experimental design (the targeted removal of a given plant group). We argue that our approach allows for a finer and more flexible statistical control than ANCOVA or multiple linear regressions. The use of a proxy for initial disturbance as a co-variate (e.g., removed plant biomass), for instance through an analysis of co-variance (ANCOVA) is only applicable in cases where the treatments induce a similar amount of disturbance. This will often not be the case as plant species or groups are not equally distributed in an ecosystem. In our case, where the different plant removal treatments target plant groups that are either dominant (on average 87% cover) or sub-dominant in the community (13%) and where the controls (0%) show no variation in the covariate, the ANCOVA assumption of homogeneous variances of the covariate between groups cannot be met. While the difference between groups may be particularly large in our mycorrhizal-removal experiment, it is common to have unequal abundances of different targeted groups (Symstad & Tilman, 2001; Bret-Harte *et al*., 2004; Robroek *et al*., 2017). The assumption of homogenous regression slope coefficient across groups also cannot be met when an undisturbed control group with no variation in the covariate is included, a common issue in similar experiments. Multiple regressions provide insights on the amount of variation shared between biomass removed and other experimental (or response) variables, but offer less power than ANCOVA and are typically used when ANCOVA is not an option (e.g.; Bret-Harte *et al*., 2004). Their usefulness in model comparisons can suffer from strong collinearity: in such cases the covariate (e.g., biomass removed) may appear as uninformative and may therefore be inadequately discarded in favor of other variables, although it may be the cause for changes in these other variables. The need for homogeneous distribution of the covariate between groups is not present in our approach, as the relationship defined in our gradient regressions covers the whole range of disturbances from undisturbed to fully disturbed. So long as the gradient encompasses the disturbance levels that the different treatments cause, it is not necessary that the treatments are similar to each other in terms of the disturbance covariate. Further, ANCOVA assumes a linear response between the response variable and the covariate. In the case where the relationship between the covariate and the response variable is linear, and if the treatments remove a similar amount of biomass, then our gradient approach is essentially analogue to an ANCOVA (with the important distinction that the response-covariate relationship is here derived from a separate set of observations). In the examples we present, singular model fits lead us to using mainly linear models, however our approach can be applied beyond linear models to non-linear relationships, such as mixed models or generalized linear (mixed) models, as exemplified.

Not adjusting for non-specific biomass removal effects in the variables we present would have led to mistaken ecological inferences. For instance, the increase in *B. nana* δ^15^N with ericoid plant removal (Fig. 4b, 4c) could have been attributed to altered competition dynamics between ericoid and ectomycorrhizal fungi. In fact, it purely results from the removal of plant biomass, and thus the observed increase might be instead related to decomposition of root necromass, while the removal of ericoid plants did not further affect *B. nana* δ^15^N content. Conversely, the non-significant effect of ericoid plant removal on *B. nana* growth appeared as a strong decrease after adjusting for non-specific biomass removal. The negative effect on growth compared to the control could for instance stem from facilitation dynamics between ericoid plants and *B. nana*. Without adjustment, the non-significant effect of ericoid removal might not have been discussed, or even reported, due to publication bias against non-significant results (Rosenthal, 1979; Marks-Anglin & Chen, 2020; Nakagawa *et al*., 2022).

## Conclusion

We present an experimental design that allows to partition non-specific disturbance effects from targeted disturbance, in the context of a mycorrhizal plant removal experiment. This approach allows us to identify and circumvent biases that can lead to mis-estimated effect sizes or plainly incorrect interpretations. Compared to the commonly used approaches of waiting for a hypothetical recovery of the system, or using statistical control tools like ANCOVA, our hybrid approach provides more possibilities to adjust response variables and control for biases induced by initial disturbance.

## Acknowledgements

We thank the Swedish Polar Research Secretariat and SITES for the logistical support of the work done at the Abisko Scientific Research Station. This study was financially supported by the Bolin Centre for Climate Research, Helge Ax:son Johnsons stiftelse and Arcum (SM); Swiss Polar Institute Exploratory Grant (SPIEG-2020-001) and Swiss National Science Foundation (grant PZ00P2_174047) (KG); Waldemar von Frenckells stiftelse, Oskar Öflunds stiftelse, the Osk. Huttunen Foundation and The University of Oulu & The Academy of Finland PROFI4 (Grant 318930) (MV).

We thank James T. Weedon for critical input on statistical data processing, Anika Mayr for obtaining and processing data on *Betula nana* growth and isotopic nitrogen content, and Sonia Meller for lending us the SEAR scanner. We also thank Anna Miettinen, Elisa Jung, Henrike Lange, Janne Welling, Lorenzo Masini, Luca Dettmers, Margaux Chadanson, Paul Schulz and Yannick Bernard for their help in the maintenance of the experiment and data collection.

## Author contributions

Sylvain Monteux, Gesche Blume-Werry, Konstantin Gavazov, Eveline J. Krab, Signe Lett, Emily Pickering Pedersen and Maria Väisänen designed and carried out the experiment.

Sylvain Monteux, Gesche Blume-Werry, Konstantin Gavazov, Leah K. Kirchhoff, Eveline J. Krab, Signe Lett, Emily Pickering Pedersen and Maria Väisänen acquired, analysed and interpreted the data. Sylvain Monteux and Maria Väisänen led the writing of the manuscript. All authors contributed critically to the drafts and gave final approval for publication.

## Data and code availability

The data and code that support the findings of this study are openly available in the Bolin Centre for Climate Research Database at http://doi.org/[doi].

\*\**the code and data are in a git repository that can be shared anonymously with reviewers upon request, the code and data will be publicly released and a DOI will be minted by the Bolin Centre Database once the manuscript is accepted, and this section will be updated subsequently*\*\*

## Supplements

**Supplementary Figure S1:**
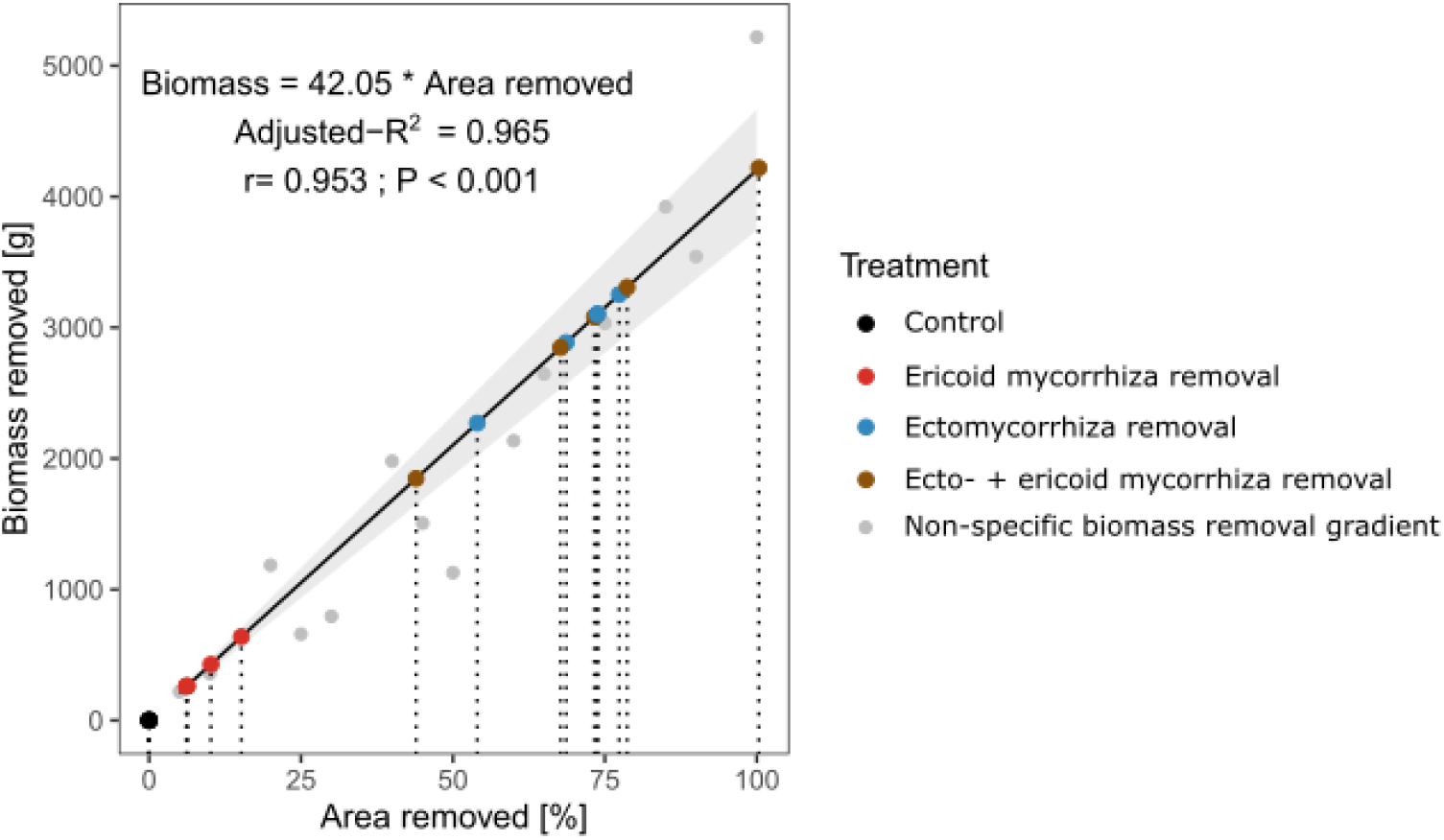
Relationship between non-specific biomass removal and percentage of area where biomass was removed. The grey symbols represent the non-specific biomass removal gradient plots, the regression line is fitted on these plots with a fixed intercept at 0, statistics denote that linear regression and the Pearson correlation test between area and removed biomass. Targeted removal treatment plots are fitted on the regression based on the amount of biomass removed to estimate the equivalent area removed in each plot.

**Supplementary Figure S2:**
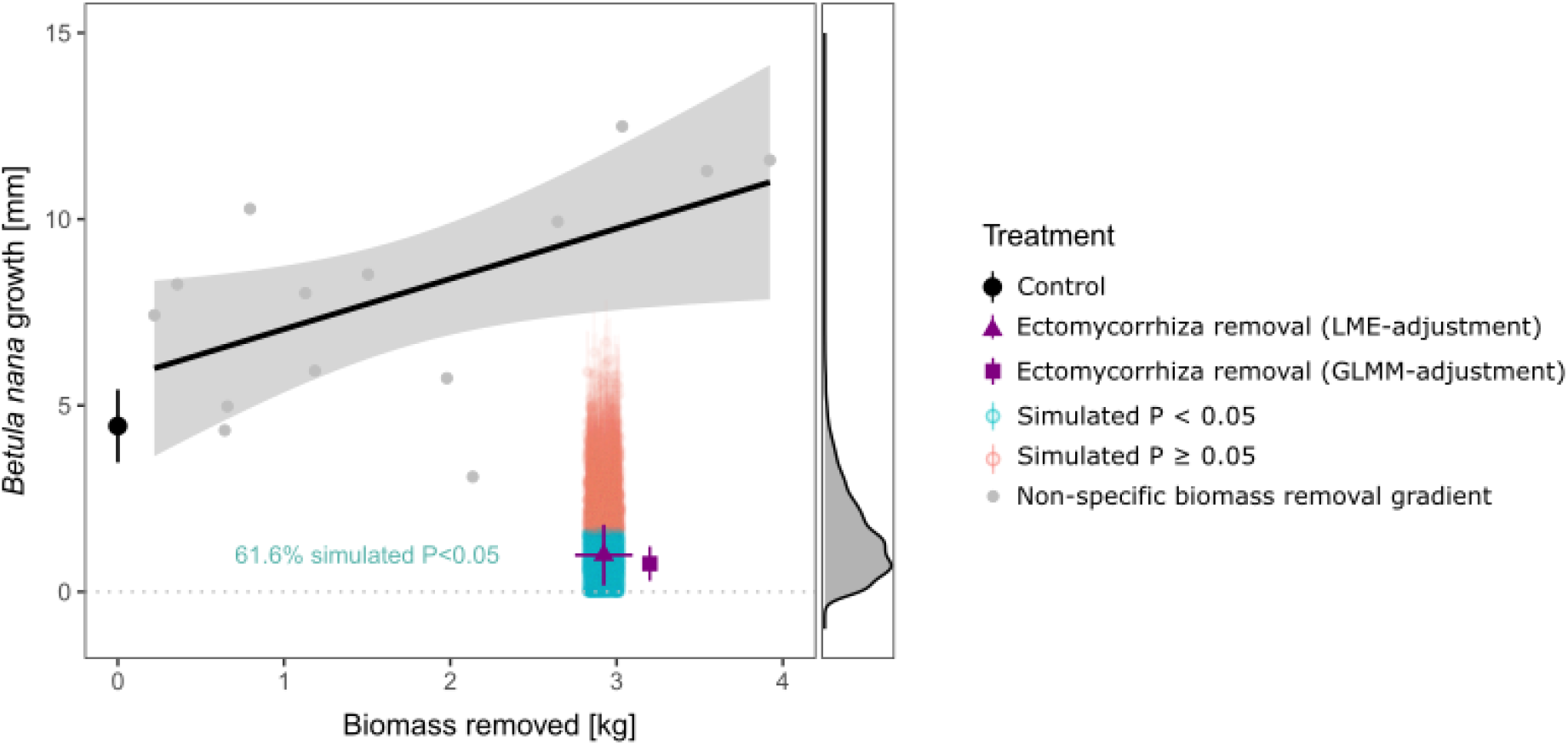
Adjusted and simulated *Betula nana* growth values when replacing negative adjusted growth values by a small positive value (10^−5^). The grey symbols are from the non-specific biomass removal gradient (grey dots), showing also the corresponding gradient régression (black line, solid when régression P < 0.05, otherwise dotted) and its 95% confidence interval (grey shaded area). The purple triangle represents the mean +/- SE (n = 5) of *Betula nana* growth after adjustment using linear mixed model (LME) and replacing negative adjusted values by 10^−5^. The blue and red transparent symbols each represent the mean +/- SE of one of the 5000 simulated datasets, respectively when the *P* value of the ANOVA test for the -ECM treatment effect is below or above 0.05. The inset on the right-hand side represents the density distribution of simulated means. The purple square represents mean +/- SE (n = 5) after adjustment using generalized linear mixed model (GLMM) with an inverse link on a Gamma distribution, also replacing negative adjusted values by 10^−5^. The simulated points and the GLMM-adjusted data have the same value on the x-axis as the LME-adjusted data and are only spread out for readability.

